# Bab2 activates JNK signaling to reprogram *Drosophila* wing disc development

**DOI:** 10.1101/2020.12.30.424794

**Authors:** Yunpo Zhao, Jianli Duan, Alexis Dziedziech, Sabrina Büttner, Ylva Engström

## Abstract

In response to cellular stress and damage, certain tissues are able to regenerate and to restore tissue homeostasis. In *Drosophila* imaginal wing discs, dying cells express mitogens that induce compensatory proliferation in the surrounding tissue. Here we report that high levels of the BTB/POZ transcription factor Bab2 in the posterior compartment of wing discs activates c-Jun N-terminal kinase (JNK) signaling and local, cell-autonomous apoptotic cell death. This in turn triggered the upregulation of the Dpp mitogen and cellular proliferation in the anterior compartment in a JNK*-*dependent manner. In the posterior compartment, however, *dpp* expression was suppressed, most likely by direct transcriptional repression by Bab2. This dual-mode of JNK-signaling, autocrine pro-apoptotic signaling and paracrine pro-proliferative signaling, led to opposite effects in the two compartments and reprogramming of the adult wing structure. We establish Bab2 as a regulator of wing disc development, with the capacity to reprogram development via JNK activation in a cell-autonomous and non-cell-autonomous manner.

**Summary statement:** Zhao et al. shows that the BTB/POZ transcription factor Bab2 is a potent activator of JNK signaling, apoptosis and compensatory proliferation, thereby driving both pro-tumorigenic and anti-tumorigenic processes.

## Introduction

Regeneration refers to the organismal processes that ensure repair and replacement of lost or damaged tissues, or promote the continuous renewal of epithelial linings. It requires cellular proliferation combined with proper cell fate specification and patterning. There are many experimental models to study regenerative growth, but until recently, these relied primarily on surgical wounding/removal/transplantation of limbs and tissues. With the development of genetic ablation of cells and tissues, alternative non-invasive techniques have become amenable in many organisms. In model organisms like zebrafish and *Drosophila*, the underlying genetic programs that initiate regeneration and cellular proliferation are being dissected in great detail. Many questions still remain though, and refined models will be important to get a comprehensive and mechanistic understanding of these processes, both during normal development as well as in response to physical stress/injury, infections, and tumorigenesis.

To prevent tissue overgrowth and ensure tissue homeostasis, regenerative growth is largely coupled to, or a result of, cell death. *Drosophila* larval imaginal discs are a favorable model for studying the coupling between cell death and regenerative growth(Sun and Irvine, 2014, Beira and Paro, 2016, Hariharan and Serras, 2017). Regeneration occurs in response to removal of part of the disc, or after induction of apoptosis by irradiation or expression of pro-apoptotic genes(Smith-Bolton et al., 2009, Sun and Irvine, 2014, Herrera et al., 2013, Grusche et al., 2011). Cells near to the dying or wounded cells proliferate in a process called compensatory cell proliferation(Fan and Bergmann, 2008, Ryoo and Bergmann, 2012).

Several signal transduction pathways are required for compensatory cell proliferation in the *Drosophila* model, such as Myc, p53, Hippo/Yorki, EGFR, and Notch signaling(Wells et al., 2006, Smith-Bolton et al., 2009, Dichtel-Danjoy et al., 2013, Sun and Irvine, 2014). In addition, the TNF/c-Jun N-terminal kinase (JNK) pathway plays a key role in response to cell stress and damage(Ryoo et al., 2004, Mattila et al., 2005, Perez-Garijo et al., 2009, Bergantinos et al., 2010, La Marca and Richardson, 2020). While JNK-activation triggers apoptosis and cell death in the cells where it is expressed, it also drives compensatory cell proliferation(Pinal et al., 2019). Furthermore, JNK activation in apoptosis-deficient *Drosophila* cells induces tumorigenesis(Perez-Garijo et al., 2009, Pinal et al., 2018). Similarly, there is substantial experimental evidence of both a tumor suppressor role of JNK in mammals, and of JNK activity coupled to cancer development and progression(Bubici and Papa, 2014, La Marca and Richardson, 2020). Thus, JNK activation can promote pro-apoptotic/anti-tumorigenic as well as regenerative/pro-tumorigenic processes in both *Drosophila* and vertebrates. An in-depth analysis of the diverse and pleiotropic effects of JNK signaling is necessary to take full advantage of this pathway as a target for for therapeutic interventions of human malignancies including cancer.

The *Drosophila bric-a-brac* locus transcription factors Bab1 and Bab2 are nuclear BTB/POZ (Broad-Complex, Tramtrack and Bric-a-brac/POZ (Pox virus and Zinc finger) domain proteins that are involved in morphogenesis, abdomen pigmentation, and development of ovaries, legs and antennae(Godt et al., 1993, Godt and Laski, 1995, Sahut-Barnola et al., 1995, Kopp et al., 2000, Couderc et al., 2002). It was previously reported that ectopic expression of *bab2* causes developmental defects in *Drosophila* imaginal tissues that are indicative of pronounced cell death while simultaneously showing signs of regenerative tissue growth(Bardot et al., 2002). In a recent study, we demonstrated that Bab2 is acting as a transcriptional repressor, downstream of the ecdysone signaling hierarchy, in regulation of *Salivary gland secretion (Sgs*) genes during the initiation of metamorphosis(Duan et al., 2020).

Here we show that the BTB/POZ transcription factor Bab2 is necessary for proper cell growth and size control of the *Drosophila* wing imaginal disc and the adult wing. Furthermore, we demonstrate that ectopic expression of *bab2* induced cell-autonomous apoptosis and non-cell-autonomous compensatory proliferation. This required both autocrine and paracrine JNK signaling, mediating upregulation of the mitogen BMP/Dpp. Importantly, Bab2 directly repressed *dpp* expression, blocking cell-autonomous compensatory proliferation. As a result, the dual functions of JNK signaling, pro-apoptotic and pro-proliferative, were mechanistically intertwined but physically separated in the two wing disc compartments. This changed the growth of the two halves of the wing disc in opposite directions, and consequently affected the adult wing structure. Thus, ectopic Bab2 expression facilitates the further analysis of JNK-driven expression of mitogenic factors and proliferation in the context of normal and malignant development.

## Results

### Bab2 is required for normal growth of imaginal discs and adult structures

Bab2 is expressed in the tarsal primordium of leg imaginal discs (Fig. 1a), in antennal discs (Fig. 1b)(Godt et al., 1993, Couderc et al., 2002), as well as in wing discs (Fig. 1c), but is absent in eye discs (Figure 1b). To analyze if Bab2 plays a role in wing imaginal disc development, *bab2* mRNA was downregulated by RNA interference (RNAi). Genetic manipulations of *bab2* and other regulatory factors in distinct regions of the wing disc was accomplished using different Gal4 driver lines (Fig. 1d). Silencing of *bab2* in the whole wing pouch region, using the *nub-Gal4* driver, reduced Bab2 protein levels considerably (Fig. 1e), indicating efficient depletion of Bab2. This decreased the final wing size (Fig. 1f,g), suggesting that *bab2* plays a role in regulation of growth, either directly or indirectly. In contrast, when *dpp-*Gal4 was used to downregulate *bab2* specifically at the A/P boundary, which constitutes a signaling center for patterning and tissue growth(Diaz-Benjumea and Cohen, 1993, Tabata et al., 1995), the wing size increased (Fig. 1h,i). Thus, silencing of *bab2* showed opposite effects on growth, depending on the site of *bab2-IR* expression.

**Fig. 1.**
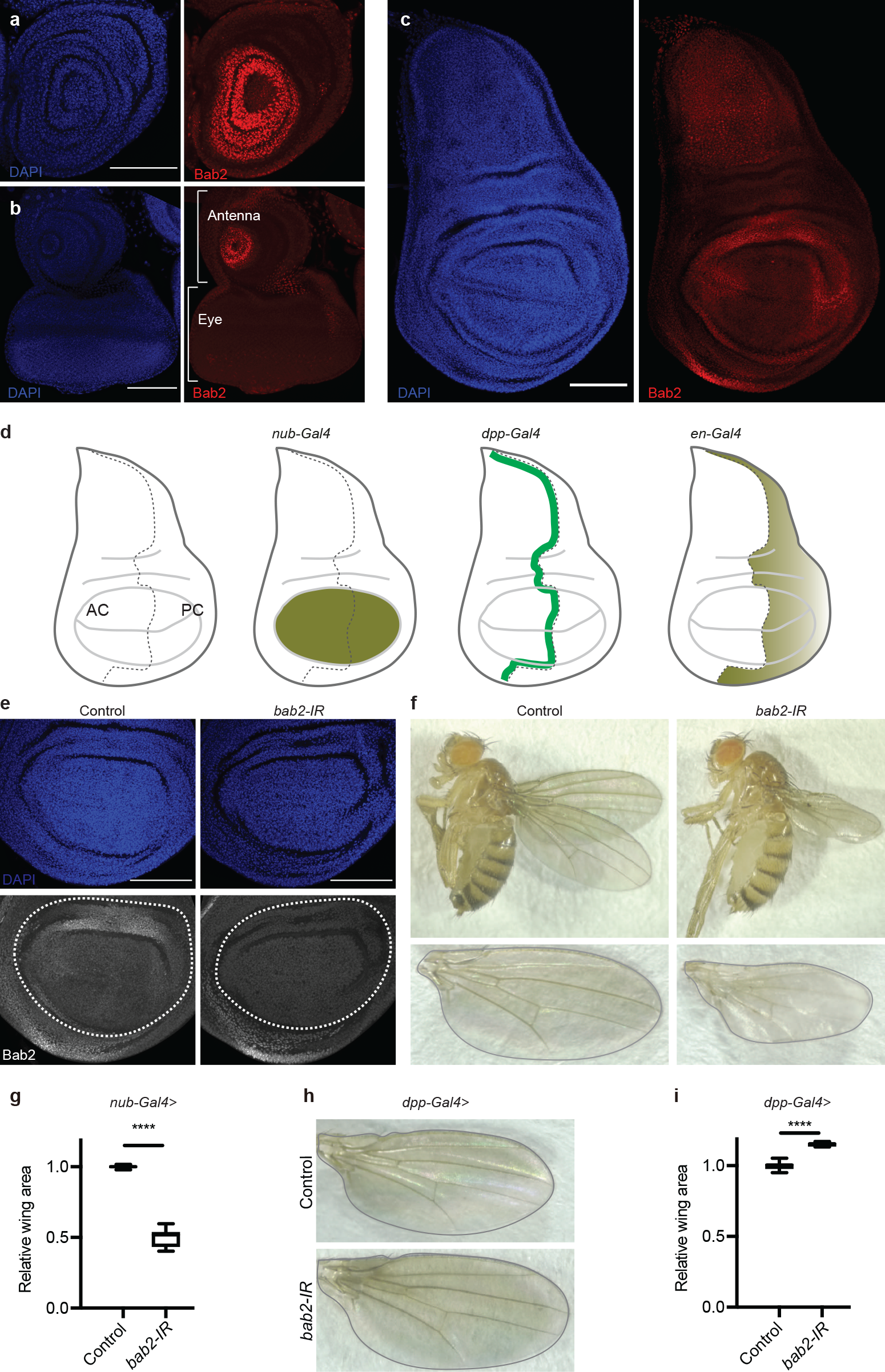
Bab2 regulates wing size. **a-c** Bab2 immunostaining and confocal fluorescence images of leg (**a**), eye-antennal (**b**), and wing (**c**) imaginal discs in *w*^*1118*^. **d** Schematic illustration of wing imaginal discs. Three different Gal4 driver lines *nub-Gal4, dpp-Gal4*, and *en-Gal4* express in distinct regions of the wing disc. AC, anterior compartment; PC, posterior compartment. **e** Bab2 immunostaining and confocal fluorescence images of wing discs after *bab2-RNAi* in the wing pouch (*UAS-Dicer2*/+; *nub-Gal4*/+) (left) and (*UAS-Dicer2*/+; *nub-Gal4*/*UAS-bab2-RNAi*) (right). Wing pouch regions are highlighted with white dashed lines in (**e**). Anti-Bab2 staining is shown in red (**a-c**) and in grey (**e**); DAPI stains DNA in blue. Scale bars represent 100 µm. **f-g** Downregulation of *bab2* in the wing pouch region reduces adult wing area. **f** Upper panels, female adult flies; lower panels, representative wings. Left, control (*UAS-Dicer2*/+; *nub-Gal4*/+); right, (*UAS-Dicer2*/+; *nub-Gal4*/*UAS-bab2-RNAi)*. **g** Quantification of adult wing area in (**f**). *n* = 6 (control) and 8 (*bab2-RNAi*). Median (middle line) is depicted and whiskers show minimum to maximum. Statistical analysis was performed using an unpaired student’s *t*-test with Welch’s correction. \*\*\*\**p*<0.0001. **h-i** Down-regulation of *bab2* in the A/P border increases wing size. **h** Upper panel, (*nub-Gal4, UAS-mCherry*/+; *dpp-lacZ*/+); lower panel, *bab2-RNAi* expression (*nub-Gal4, UAS-mCherry*/*UAS-bab2-RNAi*; *dpp-lacZ*/+). **i** Quantification of adult wing area in (**h**). *n* = 8. Median (middle line) is depicted and whiskers show minimum to maximum. Statistical analysis was performed using a *t-*test with Welch correction. **** *p*<0.0001.

As a first step in understanding how Bab2 regulates wing disc growth, we analyzed the expression of the *Drosophila* morphogens Dpp and Wg, which also act as mitogens during wing disc development(Perez-Garijo et al., 2004, Ryoo et al., 2004, Wells et al., 2006, Smith-Bolton et al., 2009, Barrio and Milan, 2017, Bosch et al., 2017, Matsuda and Affolter, 2017, Barrio and Milan, 2020). Downregulation of *bab2* increased *dpp-*driven *lacZ* expression in a region in the dorsal part of the wing pouch (Fig. S1a,b), where *bab2* normally is highly expressed (Fig 1c), suggesting that *dpp* expression was upregulated in response to *bab2* silencing. Anti-Wg protein staining on the other hand decreased in the lateral region of the *bab2-RNAi* wing pouch compared to control (Fig. S1c,d). Taken together, silencing of *bab2* affected both Dpp and Wg expression, suggesting a role for these mitogenic factors downstream of Bab2 in the regulation of wing disc growth.

### Bab2 overexpression changes wing morphology via non-cell-autonomous induction of proliferation

To further investigate the role of Bab2 in the regulation of growth, we induced ectopic *bab2* expression in the wing imaginal discs. In order to modulate the expression in a highly controlled and cell-specific manner, we took advantage of Flp recombinase to drive ectopic expression of *bab2* in cell clones, in an otherwise normal genetic background. In control wing discs, activation of Flp recombinase induced widespread somatic GFP-expressing clones, while simultaneous expression of *UAS-bab2* decreased the density of the clones (Fig. 2a), and also reduced clone area significantly (Fig. 2b), indicating that ectopic Bab2 expression negatively affects the growth or survival of these cells. This was further confirmed by expressing Bab2 in the whole wing pouch under the control of *nub-Gal4*. This decreased the wing pouch area (Fig. 2c,d) and caused pronounced reduction of the adult wing size (Fig. 2e-g), indicating that ectopic expression of *bab2* causes cell death and/or compromised cell proliferation.

**Fig. 2.**
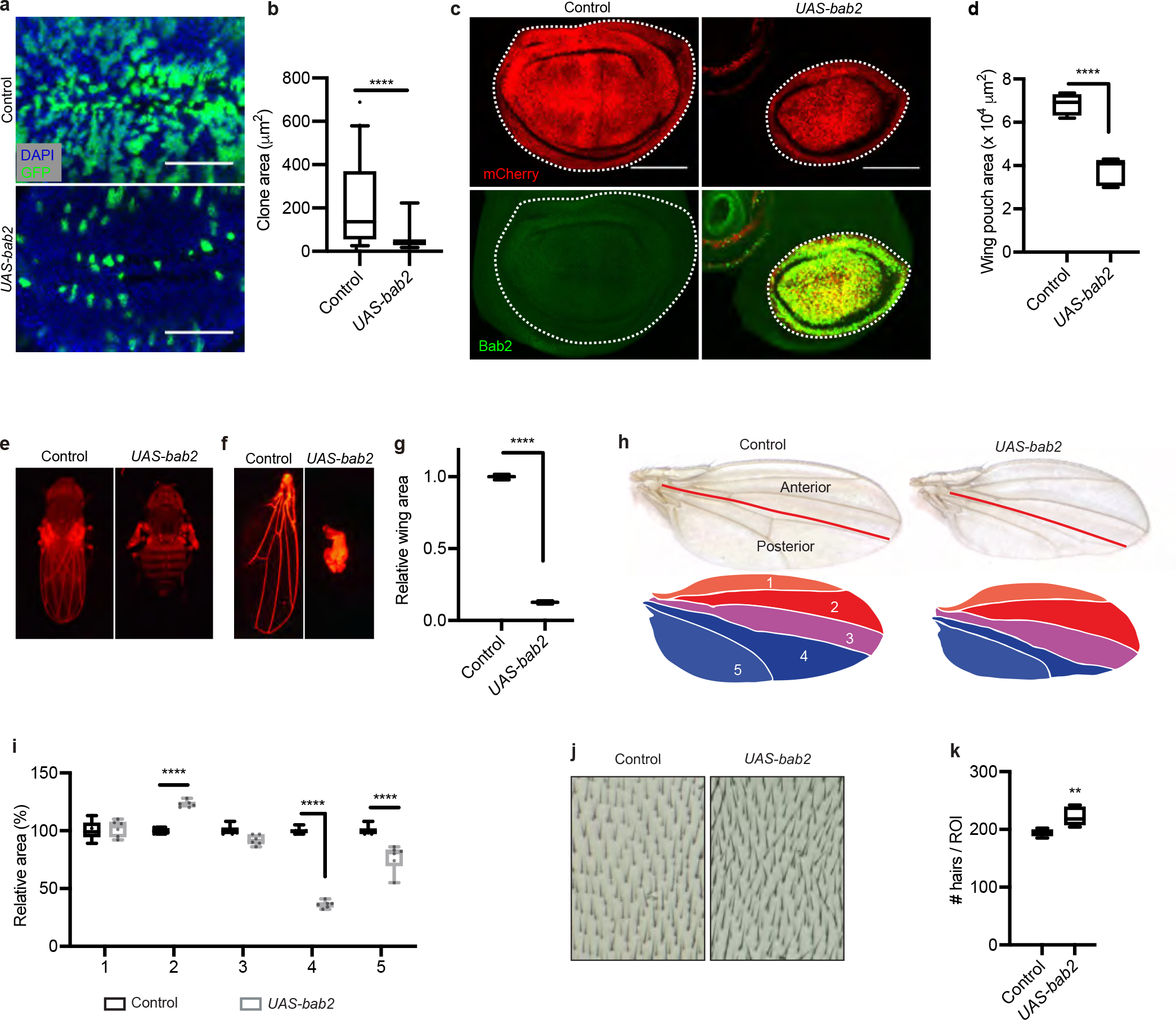
Bab2 overexpression changes wing disc growth and morphology. **a-b** Bab2 overexpression changed cell behavior in Flp-out clones in the wing discs. **a** Confocal images of wing imaginal discs (wing pouch region). Upper, control disc with *UAS-GFP*-expressing clones (shown in green); lower, wing disc with Flp-out cones expressing both *UAS-GFP* and *UAS-bab2*. Genotypes: *hs-Flp/+*; *act5C>stop>Gal4, UAS-GFP/+* and *hs-Flp/+*; *UAS-bab2/+*; *act5C>stop>Gal4, UAS-GFP/+*. Scale bars represent 50 µm. **b** Quantification of clone area of control wing discs against *UAS-bab2* overexpression discs. *n* = 40. Median (middle line) is depicted and whiskers show minimum to maximum. Statistical analysis was performed using a *t-*test with Welch correction. **** *p*<0.0001. **c-g** Overexpression of Bab2 in the wing pouch. **c** Confocal images of wing imaginal discs, mCherry labels wing pouch in red (outlined with white dashed lines); immunostaining of Bab2 (green). Control (*nub>mCherry*) and Bab2 overexpression (*nub>mCherry, bab2*). Scale bars represent 100 µm. **d** Quantification of wing pouch area in (**c**), *n* = 4 and 6. **e-f** Fluorescent images of newly eclosed adult flies (**e**) and of adult wings (**f). g** Quantification of wing area in (**f**), *n* = 6 and 4. **d, f** Median (middle line) is depicted and whiskers show minimum to maximum; Statistical analysis was performed using a *t*-test with Welch’s correction. ****, *p*<0.0001. **h-k** Overexpression of Bab2 in the PC. Control (*en-Gal4*/+) and Bab2 overexpression (*en-Gal4*/*UAS-bab2*). **h** Upper panels, wings of female adults. Anterior is up, distal to the right. The A/P boundary is indicated by a red line. Lower panels, schematic illustrations of intervein areas in female wings. The five intervein regions IV 1-5 are color-coded as light red, red, magenta, blue, and light blue. **i** Quantification of intervein areas. Flat wings (i) and flat wings with a notch (ii) (Fig S2) were used for the quantification, while crumbled wings (iii) were excluded. *bab2* overexpression in the posterior compartment significantly reduced IV4 and IV5. It also led to significant enlargement of IV2 in a non-autonomous manner. Statistical analysis was performed with a two-way ANOVA corrected with Sidak, *n* = 6. \*\*\*\**p*<0.0001. **j-k** Quantification of cell density in IV2, using the number of wing hairs in a specified wing area (region of interest, ROI) as a proxy for cell numbers. Representative images (**j**) and quantification (**k**), *n* = 6. Median is depicted and whiskers show minimum to maximum values; statistical analysis was performed using a student’s *t*-test with Welch’s correction. **p<0.01.

Next we manipulated *bab2* expression selectively in the posterior compartment (PC), using *engrailed-Gal4 (en-Gal4*), and assessed wing morphology. The *Drosophila* wing consists of single dorsal and ventral layers of epithelial cells. Five longitudinal veins, L1-L5 transverse the wing sheet and divide the wing into five interveins (IVs), designated IV1-IV5 from anterior to posterior (Fig. 2h). The *en-Gal4* driver is expressed in progenitor cells of IV4 and IV5, as well as in the posterior part of IV3. Interestingly, *en-Gal4*-driven expression of *bab2* in wing discs changed wing morphology in a consistent manner but with graded severity (Fig. 2h and Fig. S2a,b). The observed wing phenotypes ranged from (i) flat wings and (ii) flat wings with distal notches to (iii) severely wrinkled wings, each accounting for about one-third of the wings (Fig. S2a,b), suggesting a graded penetrance of aberrant wing morphology. Detailed analysis and quantification of intervein areas revealed that *bab2* expression in the PC led to a reduction of the IV4 and IV5 area, while conversely the IV2 area increased compared to interveins in control discs (Fig. 2h,i). To determine whether the enlargement of IV2 was caused by an increase in cell size or cell numbers, we quantified the cell density in IV2. Compared to control, *bab2* expression increased the number of wing hairs in IV2 (Fig. 2j,k), indicative of higher cell density. Thus, the enlargement of IV2 is likely caused by over-proliferation and not by increased cell size. In conclusion, Bab2 expression in the whole wing pouch triggers a prominent reduction of wing size. Instead, ectopic expression of Bab2 selectively in the PC of wing discs only reduced the posterior wing interveins, while it stimulated cell proliferation in distinct anterior wing primordial tissues. Importantly, this indicates that Bab2 has opposite effects on tissue maintenance and growth in the two wing disc compartments, and prompted further investigation to gain a mechanistic understanding of this phenomenon.

### Bab2 induces apoptosis cell-autonomously

To investigate if the decrease of distinct intervein areas of adult wings observed upon ectopic expression of *bab2* involved apoptotic cell death we tested for markers of apoptosis. Again, *bab2* expression was manipulated in the PC of wing discs, and the anterior compartment (AC) was used as an internal control tissue. The pro-apoptotic gene *reaper (rpr)* promotes apoptosis when Rpr and Diap1 proteins interact and release the inhibition of caspases(Goyal et al., 2000, Wilson et al., 2002). Thus, induction of *rpr*-*lacZ* serves as a read-out for the activation of apoptosis. A basal level of *rpr-lacZ* expression was observed in both the AC and PC of control discs, as revealed by immunostaining against β-galactosidase (Fig. 3a). Upon *bab2* transgene expression in the PC of wing discs, the fluorescence signal of *rpr-lacZ* increased in the PC, and weakly in the AC (Fig. 3a,b), suggesting that *bab2* overexpression induced apoptosis. Moreover, immunostaining against Cleaved Caspase 3 (CC3) revealed prominent cleavage of caspases in the PC as a result of *bab2* expression, indicative of activation of apoptosis (Fig. 3c). No apoptotic caspase cleavage was detectable in the AC (Fig. 3c,d), indicating that *bab2* expression induced apoptosis in a cell-autonomous manner. Collectively, these data suggest that PC-restricted upregulation of *bab2* expression triggers cell-autonomous apoptosis in the PC (Fig. 3a-d) and stimulates non-cell-autonomous proliferation in the AC (Fig. 2a,h-k). Moreover, it also suggests that the pronounced reduction in size of both wing discs and adult wings (Fig. 2e-g), after ectopic *bab2* expression in the whole wing pouch, is the result of activation of apoptotic cell death, possibly combined with loss of compensatory cell proliferation.

**Fig. 3.**
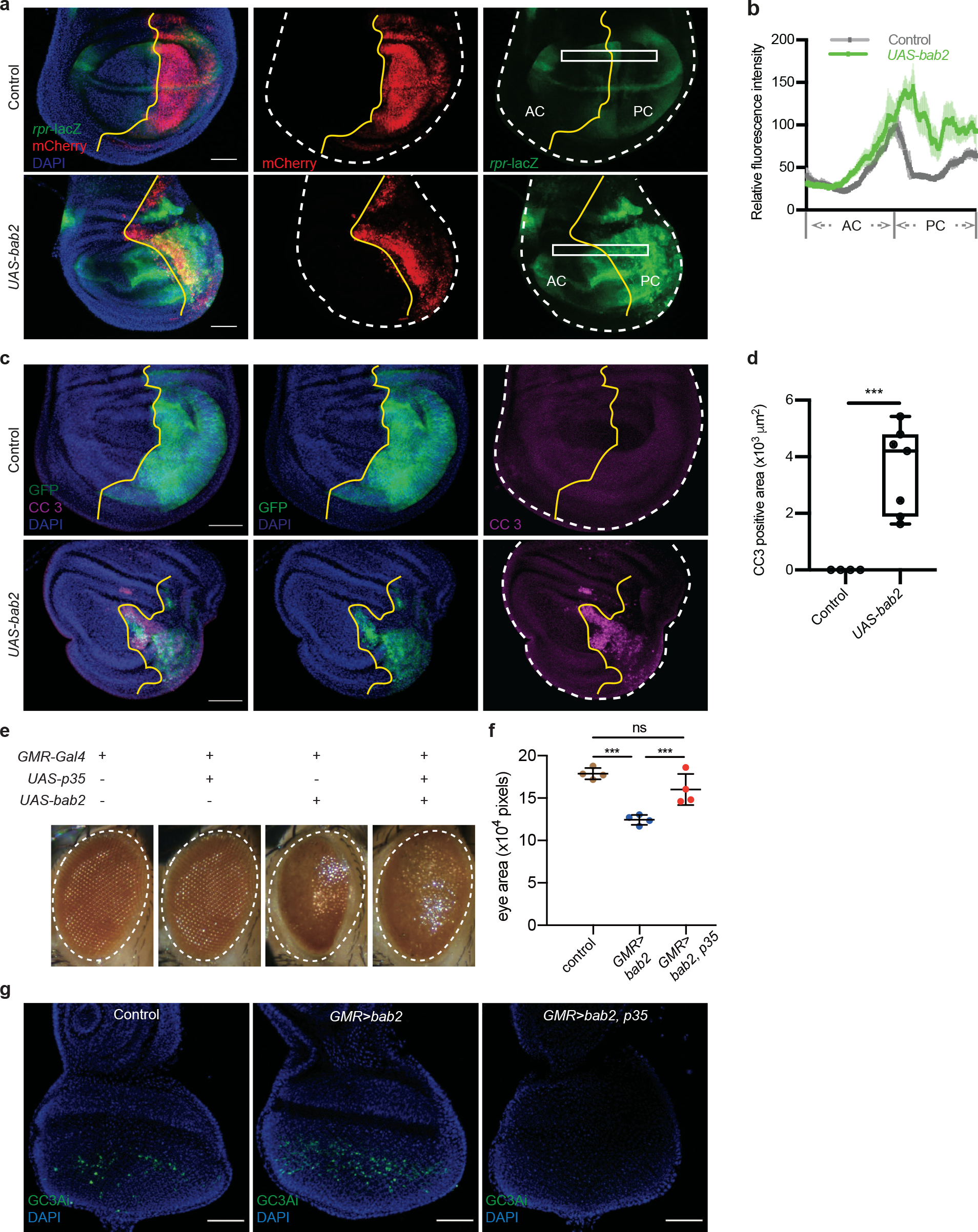
Bab2 expression induces apoptosis in imaginal discs. **a-d** Bab2 overexpression in the PC of wing discs. **a** Confocal images of wing imaginal discs, as outlined white dashed lines, with *rpr-lacZ* expression (green), the PC is marked by mCherry (red), A/P compartment boundaries are marked by yellow lines, DAPI stains DNA in blue. Upper panel, control (*en-Gal4, UAS-mCherry*/+; *rpr-lacZ*/+); lower panel, *UAS-bab2* (*en-Gal4, UAS-mCherry*/ *UAS-bab2*; *rpr-lacZ*). Scale bars represent 50 µm. **b** Plot showing the relative fluorescence intensity in boxed regions in (**a**). *n* = 3 for each genotype. **c** Wing discs stained with anti-Cleaved Caspase 3 (CC3) in magenta, the PC is marked by GFP (green), DAPI stains DNA in blue. Upper panel, control (*en-Gal4, UAS-GFP/+*); lower panel, *UAS-bab2* (*en-Gal4, UAS-GFP/UAS-bab2*). Scale bars 50 µm. **d** Quantification of apoptotic area in wing imaginal discs in (**c**), *n* = 4 and 7, respectively. Median (middle line) is depicted and whiskers show minimum to maximum. Statistical analysis was performed using a *t-*test with Welch’s correction. *** *p*<0.001. **e-g** Ectopic Bab2 expression in eye imaginal discs. **e** Representative images of female adult eyes. Control flies (*GMR-Gal4)* and *GMR-Gal4>UAS-p35* flies showed regularly arranged ommatidia, while expression of Bab2 (*GMR-Gal4>UAS-bab2)* led to rough and small eyes. This was partially rescued by co-expression of *UAS-p35*. **f** Quantification of female adult eye area. The area of *GMR-Gal4>UAS-bab2* eyes was significantly smaller than in the control. *UAS-p35* rescued the small-eye phenotype caused by *UAS-bab2* expression. Statistical analysis was performed with one-way ANOVA. \*\*\**p*<0.001. **g** *GMR-Gal4* driving the expression of the active caspase tracker *UAS-GC3Ai* in the eye discs of control (left, *GMR-Gal4>UAS-GC3Ai*), *UAS-bab2* overexpression (middle, *GMR-Gal4>UAS-GC3Ai, UAS-bab2*), and *UAS-bab2* and *UAS-p35* co-expression (right, *GMR-Gal4>UAS-GC3Ai, UAS-bab2, UAS-p35*) flies. Scale bars represent 50 µm.

To test for Bab2-dependent activation of apoptosis in another tissue, we monitored the effects of ectopic expression of *bab2* in the eye imaginal discs, where it is normally not expressed (Fig. 1b). Expression of *bab2* using a strong driver for eye imaginal discs, *GMR-Gal4*, led to a small, rough eye phenotype (Fig. 3e), implying activation of apoptosis. While the rough eye phenotype was only partially restored by simultaneous expression of the caspase inhibitor protein P35 (Fig. 3e), the *bab*2-induced size reduction of the eye area was significantly rescued by P35 expression (Fig. 3f), confirming that the smaller eye was due to *bab2-*induced apoptosis. We also analyzed apoptosis using *UAS-GC3Ai*, which serves as a GFP-based caspase 3-like activity indicator(Schott et al., 2017). A small number of eye progenitor cells showed fluorescence in control discs (Fig. 3g). Upon *bab2* overexpression, the number of cells displaying caspase activity increased, while simultaneous *p35* expression completely blocked this response (Fig. 3g), demonstrating that P35 inhibited *bab2*-induced apoptotic cell death. Thus, ectopic expression of *bab2* activated a cell-autonomous apoptotic program in epithelial cells of the wing and eye imaginal discs. Moreover, by mis-expression of *bab2* in the PC of wing discs, we were able to trigger compensatory proliferation in the AC (Fig. 2).

### Bab2-induced apoptosis requires JNK

The c-jun N-terminal kinase (JNK) signaling pathway is triggered by a variety of insults, e.g., UV treatment(Adler et al., 1995), oxidative stress(Wang et al., 2003), immune challenge(Delaney et al., 2006), and host-microbe dysbiosis(Lindberg et al., 2018, Zhou and Boutros, 2020). It is also well-documented that wounding of *Drosophila* imaginal discs leads to activation of the JNK pathway(Ryoo et al., 2004, Bosch et al., 2005, Mattila et al., 2005, Perez-Garijo et al., 2009, Bergantinos et al., 2010, Katsuyama et al., 2015, Pinal et al., 2019). To investigate if the JNK pathway is activated in response to ectopic *bab2* expression we used the tetradecanoylphorbol acetate response element-GFP (TRE-GFP) line, a JNK pathway reporter with four optimal Jun/Fos heterodimer binding sites placed upstream of a minimal promoter(Chatterjee and Bohmann, 2012). A weak basal level of GFP fluorescence was observed in control wing discs, while *bab2* expression strongly induced GFP expression in the PC and to some extent also in the AC, indicating activation of JNK signaling (Fig. 4a). To improve spatial resolution, we generated heat maps of GFP fluorescence intensity for the boxed regions in Fig. 4a, revealing that the *TRE-GFP* reporter was strongly activated in the PC upon *bab2* expression (Fig. 4b), but also weakly activated in the AC (arrow in Fig 4b). Quantification of the GFP signal confirmed the increased TRE activation in the anterior and posterior compartments (Fig. 4c), demonstrating that Bab2 expression activates JNK signaling locally and that this activation also spreads into the AC compartment.

**Fig. 4.**
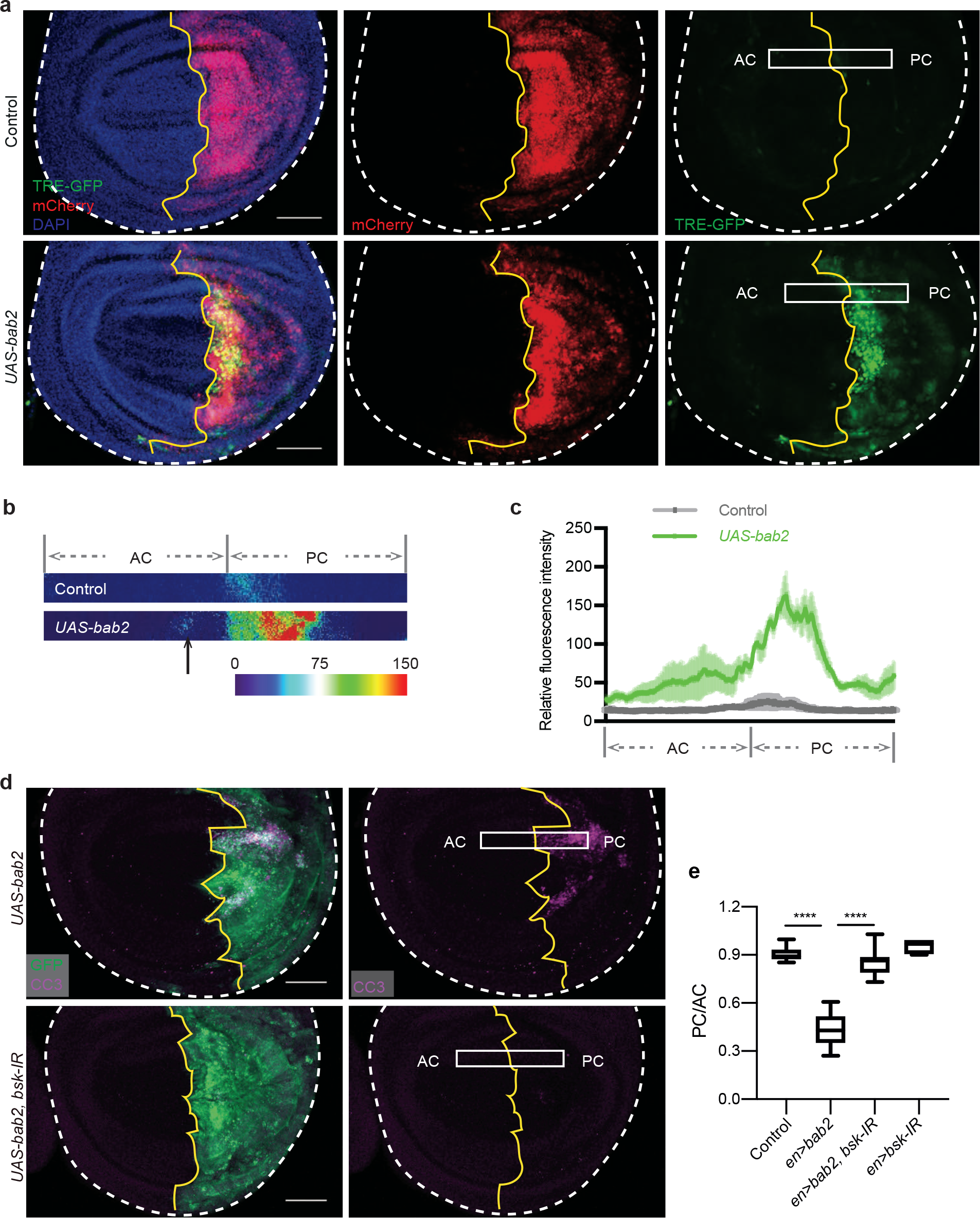
Bab2-induced apoptosis requires JNK signaling. **a-c** Overexpression of Bab2 activates JNK signaling. **a** Confocal images of wing imaginal discs, as outlined with white dashed lines, A/P compartment boundaries are marked by yellow lines. The PC is marked by mCherry (red), *TRE-GFP* expression (green), and DAPI stains nuclei in blue. Upper panel, control (*en-Gal4, UAS-mCherry*/*TRE-GFP*); lower panel, *UAS-bab2* overexpression (*en-Gal4, UAS-mCherry*/*UAS-bab2, TRE-GFP*). Scale bars represent 50 µm. **b** Heat maps represent the boxed regions in (**a**). **c** Quantification of TRE-GFP fluorescence intensity corresponding to the boxed regions in (**a**), *n* = 3 for each genotype. **d** Silencing of *bsk (JNK*) blocks Bab2-driven CC3 activation. Confocal images of wing imaginal discs, as outlined with white dashed lines, A/P compartment boundaries are marked by yellow lines. Activation of CC3 in magenta. The PC is marked by GFP (green). Upper panel, *UAS-bab2* (*en-Gal4, UAS-GFP*/*UAS-bab2*); lower panel, co-expression of *UAS-bab2* and *UAS-bsk-RNAi* (*en-Gal4, UAS-GFP*/*UAS-bab2*; *UAS-bsk-RNAi*/*+*). The CC3 signal was activated by Bab2 expression in the PC, but was abolished upon *bsk* co-expression. Scale bars represent 50 µm. **e** Plot showing the ratio of PC/AC wing disc areas in the genotypes of (**d**). Median (middle line) is depicted and whiskers show minimum to maximum; *n* = 10, 10, 7, and 6, respectively. Statistical analysis was performed with a one-way ANOVA. \*\*\*\**p*<0.0001.

JNK signaling plays a critical role in promoting apoptosis by mediating expression of pro-apoptotic genes, blocking anti-apoptotic activities, and stimulating the formation of apoptosomes and activation of caspases(Dhanasekaran and Reddy, 2017). As *bab2* expression up-regulated both JNK (Fig. 4a-c) and the apoptotic pathway (Fig. 3) in wing imaginal discs, we further investigated this link and tested whether Bab2-induced apoptosis is JNK-dependent. Upon RNAi-mediated silencing of *basket (bsk)*, encoding the *Drosophila* JNK, the Bab2-induced caspase activation in wing discs was abolished (Fig. 4d). Thus, blocking JNK expression prevented Bab2*-*induced apoptosis. We also quantified the ratio of the PC area to the AC area (PC/AC). In *en-Gal4>UAS-bab2* discs the PC/AC ratio substantially decreased compared with control discs, while co-expression of *UAS-bsk-IR* rescued the change in compartment size (Fig. 4e). Depletion of *bsk* alone did not affect the PC/AC ratio. Collectively, this indicates that JNK is necessary for Bab2-induced apoptosis in the *Drosophila* wing disc.

### Bab2 upregulates *dpp* in the wing disc in a non-cell-autonomous manner

There are two experimental models of compensatory proliferation linked to apoptosis, also termed Apoptosis-induced Proliferation (AiP)(Fan and Bergmann, 2008, Ryoo and Bergmann, 2012, Perez-Garijo, 2018). In the first, the so-called ‘genuine’ AiP model, activation of caspases triggers apoptosis and the release of mitogenic signals from the dying cells, leading to the activation of proliferation of neighboring cells(Ryoo and Bergmann, 2012, Fan et al., 2014). In this model, the non-cell-autonomous proliferation is temporally induced and compensates for cell elimination without leading to cellular overgrowth, thereby regaining homeostasis(Perez-Garijo, 2018). In the other, so-called ‘undead cell’ model, the induction of apoptosis is combined with the expression of the baculoviral protein P35, which blocks effector caspases(Perez-Garijo et al., 2004, Ryoo et al., 2004, Huh et al., 2004, Kondo et al., 2006, Fan and Bergmann, 2008, Martin et al., 2009, Perez-Garijo et al., 2009). Thus, the execution of apoptosis is inhibited in the P35-expressing cells, and these are kept alive in an artificial manner and referred to as ‘undead’ cells(Perez-Garijo, 2018). Activation of apoptosis in both these models causes mis-expression and secretion of the mitogens Wg and Dpp, promoting compensatory proliferation of neighboring cells(Perez-Garijo et al., 2004, Ryoo et al., 2004, Wells et al., 2006, Smith-Bolton et al., 2009). To unravel the molecular mechanism underlying the enlargement of the wing IV2, triggered by *bab2* overexpression in the PC, we analyzed the expression of Wg and Dpp in wing discs. Surprisingly, Wg immunostaining was reduced in both the PC and AC of wing discs upon *bab2* expression in the PC (Fig. S3a,b). The reduction of Wg staining in the AC was not expected, as *bab2* was specifically overexpressed in the PC. This result was further confirmed by immunostaining against Armadillo (Arm), a *Drosophila* homolog of β-catenin and a key player in Wg signaling(Cox et al., 1999), demonstrating reduced abundance of Arm in both the PC and AC upon *en-Gal4*-driven expression of *bab2* (Fig. S3c,d). This indicates that Wg signaling is compromised as a result of *bab2* overexpression. However, we had previously found that protein levels of Wg were reduced in parts of the wing disc upon *bab2* depletion via RNAi (Fig S1c,d). This suggests that indirect regulatory mechanisms are involved downstream of *bab2* mis-expression in the PC, which subsequently promote downregulation of Wg and Arm in the AC. Even though the mechanism of Wg downregulation in the AC is not clear, this result renders it unlikely that Wg signaling mediates the cell proliferation and subsequent IV2 enlargement in the AC, since Wg was virtually lost upon *bab2* overexpression.

The wing disc A/P compartment boundary constitutes a signaling center from which A-P patterning originates and *dpp* is expressed in a narrow strip of cells in the AC along the A/P boundary (Fig 1d)(Masucci et al., 1990). Secretion of Dpp from these cells creates an activity gradient medially to laterally, which regulates patterning and growth of the wing disc(Rogulja and Irvine, 2005, Schwank et al., 2008, Barrio and Milan, 2017, Bosch et al., 2017, Matsuda and Affolter, 2017, Barrio and Milan, 2020). To analyze whether *dpp* expression was affected by *bab2* overexpression, we again used the *dpp-lacZ* reporter. There was no overlap between the *dpp-lacZ* reporter and the mCherry-marked PC in control discs (Fig. 5a), showing that *dpp* expression was confined to cells in the AC, in accordance with previous findings(Masucci et al., 1990). However, *dpp-lacZ* strongly increased in the AC in *en-Gal4>bab2* discs (Fig. 5a,b), indicating that *dpp* was upregulated in the AC in response to Bab2 expression in the PC. As described above, Bab2 expression in the PC also led to a decrease in the ratio between PC/AC areas (Fig. 4e). We further used the *nub-Gal4* wing pouch driver to modulate *dpp* expression. In line with previous results(Barrio and Milan, 2017), RNAi-mediated downregulation of *dpp* drastically reduced wing area (Fig. 5c,d). These results are in agreement with a model in which overexpression of Bab2 in the PC leads to apoptosis-dependent upregulation of *dpp* in the AC, which then promotes compensatory cell proliferation and causes the enlargement of the IV2 wing area. It was surprising, however, that *dpp-lacZ* expression was restricted to the AC compartment as it has previously been shown that the activation of apoptosis in so-called ‘undead cells’ leads to up-regulation of *dpp* cell-autonomously in the PC(Perez-Garijo et al., 2004, Ryoo et al., 2004).

**Fig. 5.**
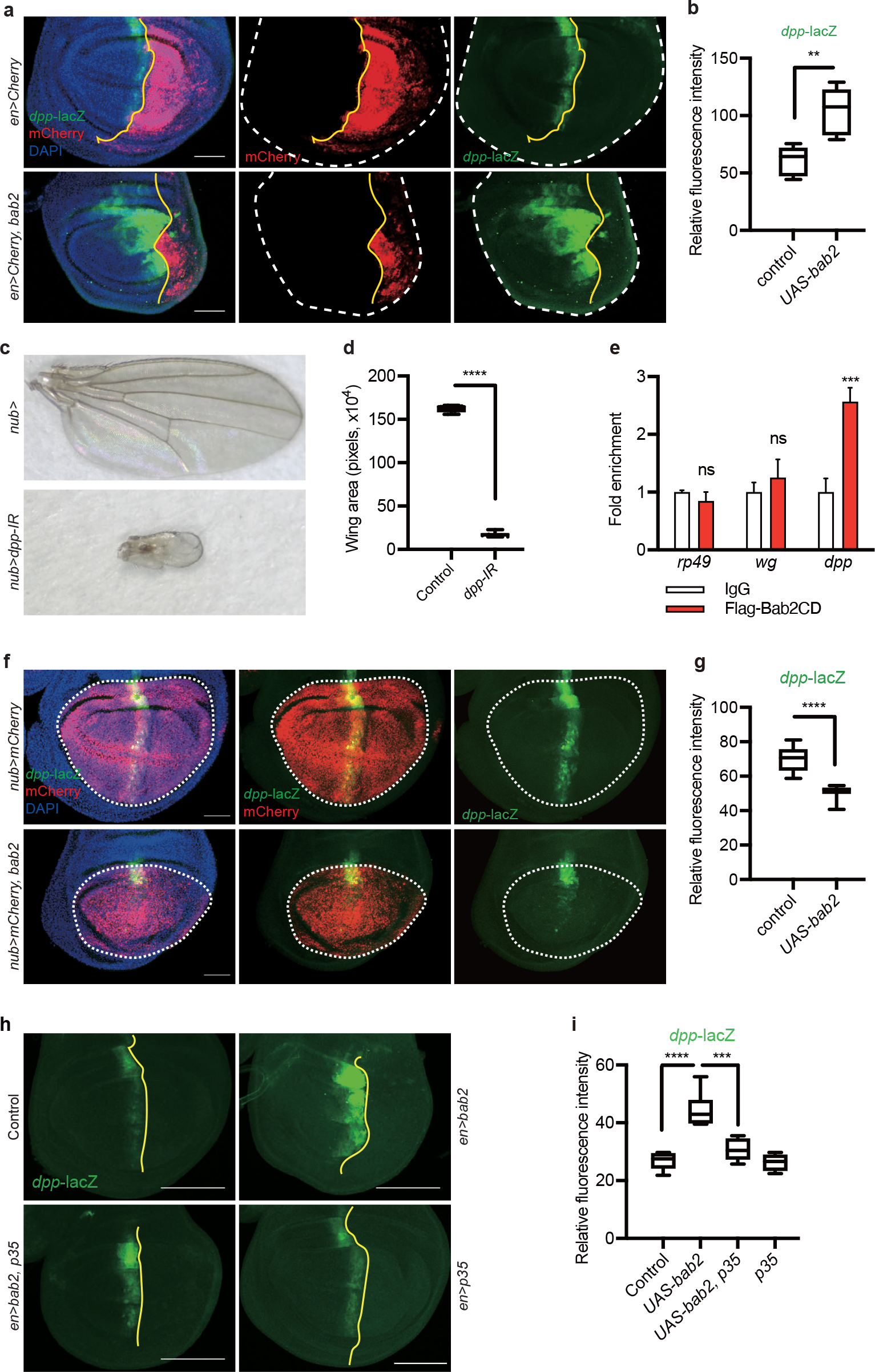
Upregulation of *dpp* in the AC by *bab2* expression in the PC, involves activation of apoptosis. **a-b** Overexpression of Bab2 in the PC and analysis of *dpp-lacZ* expression. **a** Confocal images of wing imaginal discs, as outlined with white dashed lines, A/P compartment boundaries are marked by yellow lines. The PC is marked by mCherry (red) and expression of *dpp-lacZ (*green). Upper panel, control (*en-Gal4, UAS-mCherry*/+; *dpp-lacZ*/+); Lower panel, *UAS-bab2* overexpression (*en-Gal4, UAS-mCherry*/*UAS-bab2*; *dpp-lacZ*/+). **b** Quantification of *dpp-lacZ* fluorescence intensity of the AC in the wing pouch in (**a**). *n* = 5. Median (middle line) is depicted and whiskers show minimum to maximum. Statistical analysis was performed using a *t*-test with Welch’s correction. **, *p*<0.01.) **c-d** Downregulation of *dpp* in the wing pouch led to wing size reduction. **c** Upper panel, control wing *nub-Gal4*>+; lower panel, *nub-Gal4*>*UAS-dpp-RNAi*. **d** Quantification of wing size in (**c**). *n* = 6. Median (middle line) is depicted and whiskers show minimum to maximum. Statistical analysis was performed using a *t-*test with Welch’s correction. ****, *p*<0.0001. **e** ChIP-qPCR of recombinant Flag-Bab2CD protein in S2 cells. Flag-Bab2CD is recruited to the *dpp* locus, but not to *rp49* or *wg* loci. Data represent mean with SEM. *n* = 3. Statistical analysis was performed using a one-way ANOVA with Sidak correction. ***, *p*<0.001. **f-g** Overexpression of Bab2 in the wing pouch repressed *dpp-lacZ*. **f** Confocal images of wing imaginal discs, expression of *dpp-lacZ (*green), the wing pouch is outlined by white dashed lines. Upper panel, control. *(nub-Gal4, UAS-mCherry*/+; *dpp-lacZ*/+); Lower panel, *bab2* overexpression, (*nub-Gal4, UAS-mCherry*/*UAS-bab2*; *dpp-lacZ*/+). **g** Quantification of of *dpp-lacZ* fluorescence intensity in (**f**). Median (middle line) is depicted and whiskers show minimum to maximum; *n* = 8 and 12. Statistical analysis was performed using a *t*-test with Welch’s correction. ****, *p*<0.0001. For confocal images (**a, f**), mCherry shows labeling of Gal4 expressing regions in red; DAPI stains nuclei in blue. Scale bars represent 50 µm. **h-i** Overexpression of Bab2 together with P35 in the PC blocks *dpp-lacZ* upregulation in the AC. **h** Confocal images of wing imaginal discs. The anterior of the wing discs is to the left, anterior/posterior compartment boundaries are marked by yellow lines, expression of *dpp-lacZ* (green). Upper panel, left: control (*en-Gal4, UAS-mCherry*/+; *dpp-lacZ*/+); right: *UAS-bab2* (*en-Gal4, UAS-mCherry*/*UAS-bab2*; *dpp-lacZ*/+); Lower panel, left: *UAS-bab2, p35* (*en-Gal4, UAS-mCherry*/*UAS-bab2, UAS-p35*; *dpp-lacZ*/+); right *UAS-p35* (*en-Gal4, UAS-mCherry*/*UAS-p35*; *dpp-lacZ*/+). DAPI stains DNA in blue. Scale bars represent 100 µm. **i** Quantification of *dpp*-lacZ expression in (**h)**. Median (middle line) is depicted and whiskers show minimum to maximum; *n* = 5, 6, 5, and 4, respectively. Statistical analysis was performed using a one-way ANOVA with Tukey correction. ***, *p*<0.001; ****, *p*<0.0001.

To further investigate this unexpected result, we tested whether *dpp* is a direct target of Bab2. Previous work has shown that Bab2 acts as a transcriptional repressor(Jeong et al., 2006, Gibert et al., 2007, Roeske et al., 2018, De Castro et al., 2018, Duan et al., 2020). To analyze if Bab2 binds directly to *dpp* and *wg* DNA sequence elements we performed Chromatin Immunoprecipitation (ChIP) assays, using *Drosophila* S2 cells transfected with a Flag-tagged Bab2 conserved domain (Flag-Bab2CD), containing a composite DNA binding domain(Couderc et al., 2002). Quantitative PCR (qPCR) showed that the recombinant Bab2 protein was recruited to the *dpp* locus, indicating the direct binding of Bab2 to regulatory elements of the *dpp* gene (Fig. 5e). On the contrary, there was no recruitment to the *wg* locus, confirming that the *wg* gene is not a direct target of Bab2 regulation.

Next, we investigated the effects of *bab2* expression in the whole wing pouch on *dpp-lacZ* expression. Interestingly, the normal expression pattern of *dpp-lacZ* along the A/P boundary was suppressed in most of the disc, and only a small patch of cells in the dorsal part still expressed *dpp-lacZ (*Fig 5f,g), demonstrating that Bab2 represses *dpp* expression.

Collectively, these results suggest that high-level expression of Bab2 directly repressed *dpp* and also caused cell-autonomous activation of apoptosis and JNK signaling in the PC, which in turn promoted non-cell autonomous *dpp* expression and proliferation in the AC. To assess if expression of *dpp-lacZ* in the AC, in response to *bab2* overexpression in the PC, was dependent on the induction of apoptosis, we co-expressed P35 to block cell death. Indeed, simultaneous expression of P35 inhibited the upregulation of *dpp-lacZ* in response to *bab2* overexpression (Fig. 5h,i). P35 expression *per se* had no effect on *dpp-lacZ* expression. Thus, overexpression of Bab2 represses *dpp* expression and activates apoptosis cell-autonomously, while the latter leads to *dpp* expression in adjacent cells in a non-cell-autonomous manner.

### Non-cell-autonomous *dpp* upregulation upon *bab2* expression is mediated by JNK signaling

It is well established that JNK is involved in compensatory cell proliferation downstream of apoptosis(Ryoo et al., 2004, Uhlirova et al., 2005, Bergantinos et al., 2010, Suissa et al., 2011). We next examined whether the JNK pathway plays a direct role in *bab2*-induced non-cell-autonomous *dpp* upregulation in the wing disc. To this end, we inactivated JNK signaling in the PC via RNAi-mediated silencing of the mRNA for the *Drosophila* JNK gene *basket (bsk)* and assessed Bab2-induced expression of the *dpp-lacZ* reporter. Again, overexpression of Bab2 in the PC strongly upregulated *dpp-lacZ* in the AC (Fig. 6a,b). While downregulation of *bsk* itself did not affect the normal strip of *dpp-lacZ* expression, it almost completely abolished the non-cell-autonomous *dpp* expression induced by high levels of Bab2 (Fig. 6a,b). This demonstrates that upregulation of *dpp* in the AC by Bab2 overexpression in the PC requires JNK activation and dissemination of a signal from the AC to the PC.

**Fig. 6.**
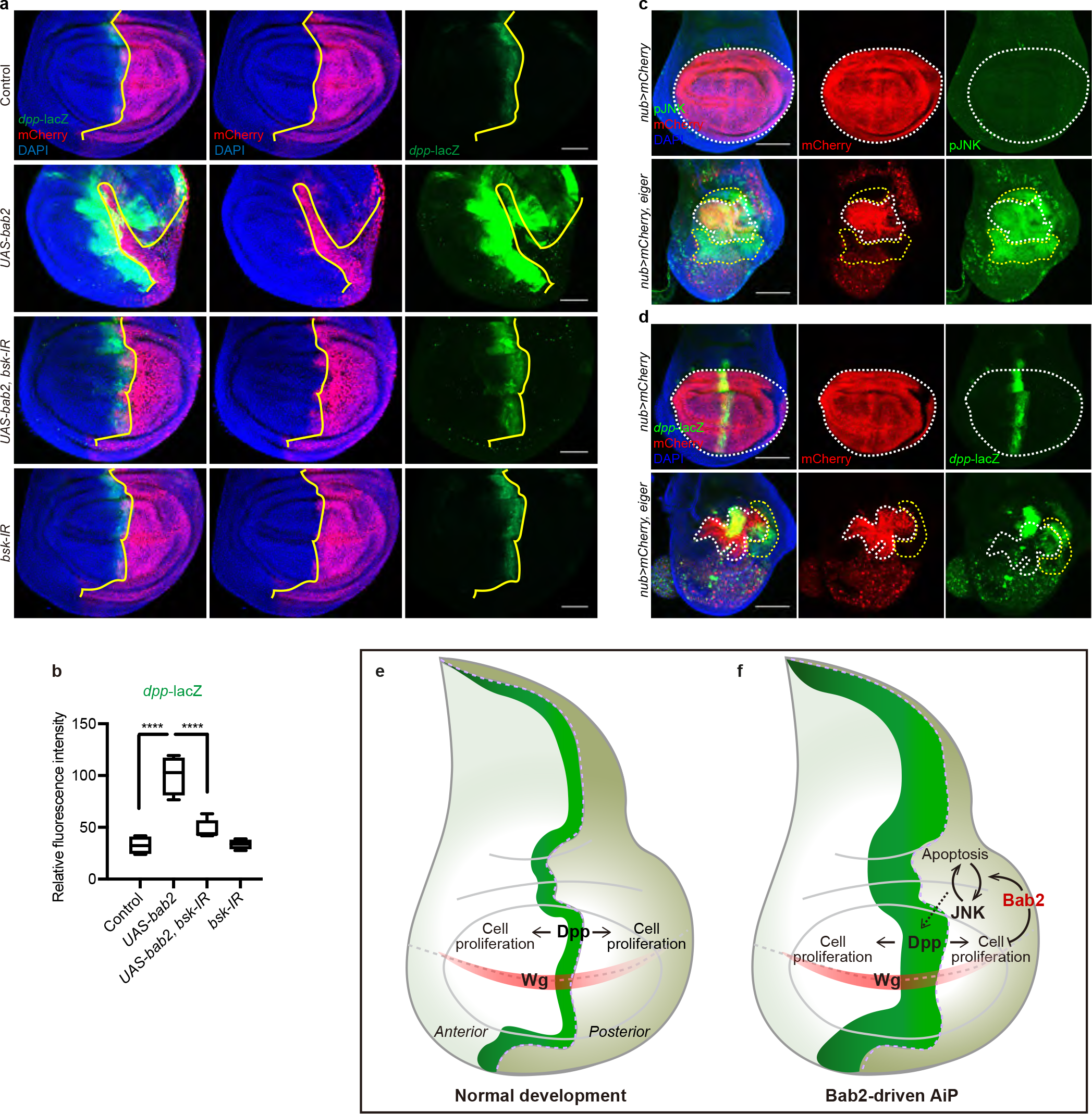
Bab2-dependent *dpp* upregulation is mediated by JNK signaling. **a-b** Overexpression of Bab2 and RNAi of *bsk (*JNK). **a** Confocal images of wing imaginal discs, A/P compartment boundaries are marked by yellow lines, the PC is marked by mCherry (red), expression of *dpp-lacZ (*green), DAPI stains nuclei in blue. The genotypes are: control (*en-Gal4, UAS-mCherry*/+; *dpp-lacZ*/+); *UAS-bab2* (*en-Gal4, UAS-mCherry*/*UAS-bab2*; *dpp-lacZ*/+); *UAS-bab2, bsk-IR* (*en-Gal4, UAS-mCherry*/*UAS-bab2*; *dpp-lacZ*/*UAS-bsk-RNAi*); and *bsk-IR* (*en-Gal4, UAS-mCherry*/*+*; *dpp-lacZ*/ *UAS-bsk-RNAi*). Scale bars represent 50 µm. **b** Quantification of *dpp*-lacZ expression (**a**). Median (middle line) is depicted and whiskers show minimum to maximum; *n* = 4, 4, 5, and 4, respectively. Statistical analysis was performed with a one-way ANOVA; \*\*\*\**p*<0.0001. **c-d** Expression of pJNK (**c**) and *dpp-lacZ* (**d**) are activated by *egr* expression. Ectopic expression of *egr* also reduces the size of the wing pouch region. Confocal images of wing imaginal discs. mCherry labels the wing pouch in red; DAPI stains nuclei in blue. **c** Activation of JNK signaling, analyzed by pJNK immunostaining (green) in cell-autonomous regions (marked with white dashed lines) and in non-cell-autonomous regions (marked with yellow dashed lines). Upper panel, control (*nub-Gal4, UAS-mCherry/+*); lower panel, *UAS-egr* overexpression in wing pouch (*nub-Gal4, UAS-mCherry/UAS-egr*). **d** *dpp-lacZ* expression (green), in cell-autonomous regions (marked with white dashed lines) and in non-cell-autonomous regions (marked with yellow dashed lines). Upper panel, control (*nub-Gal4, UAS-mCherry/+*; *dpp-lacZ/+*); lower panel, *UAS-egr* overexpression in wing pouch (*nub-Gal4, UAS-mCherry/UAS-egr*; *dpp-lacZ/+*). Scale bars represent 100 µm. **e-f** Bab2-driven AiP model. Schematic illustration of a wing disc under normal development (**e**) and with Bab2 overexpression in the PC (*en>bab2)* (**f**). The *dpp*-expressing regions are shown in bright green. The A/P boundaries are marked with dashed lines. *bab2* overexpression in the PC imposed cellular stress, which activated a loop of JNK activation and apoptosis, and decreased the size of the PC in discs, and of the IV4 and IV5 wing areas of adult wings. Direct repression of *dpp* by Bab2 in the PC blocked compensatory cell proliferation in this compartment. JNK activation in the PC promoted upregulation of *dpp* expression in the AC, which caused enlargement of the AC in discs, and the IV2 wing area.

To test whether activation of the JNK signaling would be sufficient to promote non-cell-autonomous dpp expression, we stimulated the pathway directly by expression of Eiger (Egr), a TNF superfamily ligand that activates the *Drosophila* JNK pathway. Overexpression of *egr* in the wing pouch activated JNK signaling, as measured by immunostaining against the phosphorylated from of JNK (pJNK), and drastically decreased the wing pouch area in comparison with control discs (Fig. 6c), confirming that activation of JNK signaling in the wing pouch triggers cell death, as previously reported(Smith-Bolton et al., 2009). Importantly, pJNK was also observed in regions that did not show *nub-Gal4>mCherry* expression, indicating that the JNK activation spread non-cell-autonomously (yellow outlines in Fig 6c), comparable to what we have observed using the TRE reporter (Fig 4b,c). Moreover, activation of JNK signaling via high-level expression of *egr* triggered induction of the *dpp-lacZ* reporter (Fig 6d), both inside (white outline, Fig 6d) and outside (yellow outline, Fig 6d) of *nub-Gal4* expressing regions. The pronounced depletion of cells within the wing pouch rendered the interpretation within this region difficult. It was clear, however, that *egr* overexpression within the wing pouch activated both pJNK and *dpp-lacZ* outside of the wing pouch (Fig 6c,d), demonstrating non-cell-autonomous spreading of the signal. In conclusion, activation of the JNK pathway, either by *egr* or *bab2* overexpression, triggered both cell-autonomous cell death, as well as non-cell-autonomous *dpp* expression.

## Discussion

In this study, we show that the *Drosophila* BTB/POZ transcription factor Bab2 can induce cell-autonomous apoptosis and non-cell-autonomous compensatory proliferation, leading to a reprogramming of wing disc development. Depletion of Bab2 in the A/P signaling center at the boundary between the AC and PC induced *dpp* expression and increased adult wing size (Fig 1g,h), possibly reflecting de-repression of the Dpp mitogen. *Vice versa*, overexpression of Bab2 almost abolished *dpp* expression and drastically reduced the size of wing discs, adult wings and eyes (Fig 2e-g and 3e,f). ChIP experiments confirmed that the *dpp* locus is a direct target of Bab2 binding (Fig 5e), indicating that Bab2 is a transcriptional repressor of *dpp*. On the contrary, our results do not support direct regulation of *wg* by Bab2. We conclude that Bab2 controls imaginal disc growth via direct regulation of *dpp* and most likely other growth-regulating genes and processes.

We show that ectopic expression of *bab2* in *Drosophila* wing imaginal discs leads to apoptotic cell death, and that this promotes compensatory cell proliferation in neighboring cells. The induction of apoptosis required local/autocrine JNK signaling, while the compensatory proliferation was dependent on paracrine JNK signaling and on upregulation of *dpp* in the AC of the wing disc. We propose a model in which *bab2* expression in the PC reprograms wing disc development (Fig. 6e,f). During normal development of the *Drosophila* wing disc, the expression of *dpp* in a strip of cells at the A/P boundary is necessary for normal growth and patterning(Barrio and Milan, 2017, Bosch et al., 2017, Matsuda and Affolter, 2017, Barrio and Milan, 2020). Dpp diffuses bilaterally in the wing disc, regulating cell proliferation and tissue size (Fig. 6e). Induction of apoptosis in the PC by Bab2 overexpression upregulated *dpp* non-cell-autonomously in the AC (Fig. 6f). Blocking *bab2*-induced cell death by silencing *bsk* or expressing P35 abolished *dpp* upregulation, confirming that the proliferation is indeed apoptosis-dependent and thus represents a novel mode of AiP. The Bab2-driven AiP resulted in wing disc reprogramming in the AC, which was, at least in part, mediated by the JNK and Dpp signaling pathways (Fig. 6). The apoptotic stress in the PC caused a decrease in the PC/AC ratio, and as a consequence, a reduction of the IV4 and IV5 territories of adult wings, while the IV2 territory increased as a result of over-proliferation. This indicates that apoptosis and JNK activation induced by Bab2 is constantly active. This is in agreement with a recent study, showing that transient activation of JNK causes persistent JNK function, and can lead to activation of tumorigenesis in apoptosis-deficient cells(Pinal et al., 2018). We suggest, as outlined in our model (Fig 6f), that Bab2 expression initiates a feed-back amplification loop of JNK activation and apoptosis, which subsequently causes the non-cell-autonomous upregulation of *dpp* and tissue growth in the AC. Such amplification loops, involving JNK signaling, have been described previously in apoptotic and “undead” cells downstream of different types of cellular stress, such as heat-shock, irradiation, and production of reactive oxygen species (ROS)(Shlevkov and Morata, 2012, Pinal et al., 2018, Pinal et al., 2019). Furthermore, self-reinforcement mechanisms, involving JNK signaling, secreted factors, and communication between tumor cells and surrounding tissues, drive tumor initiation and growth in several *Drosophila* models(Wells et al., 2006, Muzzopappa et al., 2017, Perez et al., 2017, La Marca and Richardson, 2020).

Importantly, unlike non-cell-autonomous proliferation driven by so-called “undead” cells in AiP, *bab2-*promoted proliferation did not involve inhibition of apoptosis by P35. Rather, it was dependent on apoptosis and continuous JNK signaling, as *dpp* expression was blocked by P35. Thus, the Bab2-driven AiP described here is P35-independent, mitogen/Dpp-dependent, and persists throughout development since it changed the final size of the wings. It thus represents an alternative AiP model, in which the molecular mechanisms downstream of apoptosis and JNK signaling are physically separated and can be investigated in the context of normal development. Considering the complex roles of JNK signaling, promoting both pro- and anti-tumorigenic effects in flies as well as mammals, experimental systems that allow a mechanistic dissection of these processes in both normal and malignant context are needed. Bab2 overexpression constitutes an alternative way to induce JNK activation in a controlled and tissue-specific manner, enabling analysis of both pro- and anti-tumorigenic functions in cells and tissues simultaneously.

The non-cell-autonomous upregulation of *dpp* implies active crosstalk between the two wing disc compartments, most likely by secreted molecules, but it may also be possible that cells migrate or change their compartment identity. In fact, when *UAS-egr* was overexpressed in the wing pouch, pJNK-positive cells were found outside of the wing pouch region (Fig 6c). Thus, it is feasible that cells with upregulated JNK activity migrate outside of the wing pouch, and that those same JNK-activated cells induce *dpp* expression. Herrara and Morata (2014)(2014) reported that genetic ablation of PC wing disc cells leads to a transient loss of the AC/PC compartment boundary, and that those cells get reprogrammed during the regeneration process in a JNK-dependent manner. In the present work, Bab2 expression in the PC triggered JNK activation and apoptosis. However, as *dpp* expression was repressed in the PC cells, it seems unlikely that those cells would become *dpp*-expressing AC cells. Therefore, we suggest that secreted molecule (s) are responsible for the cross-talk and activation of *dpp* expression and subsequent compensatory proliferation in the AC cells. Using the experimental model of compensatory proliferation described here, Bab2-driven AiP, the identification of such molecule (s) will be an important task for future investigation. This could have large implications in the understanding of tissue regeneration, and may open up new routes for JNK-based therapies of human disorders and malignancies.

## Materials and methods

### Fly Strains and genetics

Flies were maintained under a 12-h light/dark cycle and fed with a potato agar mash (12.9 g of dry yeast (Kron Jäst; Sollentuna, Sweden), 40 g of potato powder (Felix Potatismos; Skåne, Sweden), 10 g of agar (USBiological; Salem, MA USA), 50 mL of light syrup (Dansukker; Malmö, Sweden), 8.5 mL of nipagin (10% in 99% Ethanol, Sigma-Aldrich), 1 g of L-ascorbic acid sodium salt (AlfaAesar; Kandel, Germany), 4.5 mL of propionic acid, 99% pure (Acrōs Organics; Fisher Scientific), and water up to 1 L). The following fly stocks were used in the study: *w*^*1118*^ (Bloomington Drosophila Stock Center, BDSC_3605), *UAS-bsk-RNAi (*BDSC_31323), *UAS-mCherry* (BDSC_38425), *nub-Gal4, UAS-mCherry; UAS-Nslmb-vhhGFP4/TM3, sb* (BDSC_38418), *UAS-dicer2; nub-Gal4* (BDSC_25754), *UAS-egr-RNAi* (BDSC_58993), *UAS-dpp-RNAi* (BDSC_31531), *UAS-bsk-RNAi* (BDSC_31323), *rpr-lacZ* (BDSC_58793), *tub-Gal80*^*ts*^ (BDSC_7019), *dpp-Gal4* (BDSC_7007), *dpp-lacZ*(Basler and Struhl, 1994), *en-Gal4, UAS-mCD8-GFP (*provided by Adrea Brand), *UAS-bab2*(Bardot et al., 2002), *TRE-GFP*(Chatterjee and Bohmann, 2012), and *UAS-GC3Ai(Schott et al*., *2017), hs-Flp*^*122*^; *UAS-Flp*^*JD1*^*/CyO, Act-GFP*^*JMR1*^; *Act5C > CD2 > Gal4*^*S*^, *UAS-mCD8GFP*^*LL6*^*/TM6b*(Zhao et al., 2015).

To create *en-Gal4, UAS-mCherry* recombinant chromosome, *en-Gal4, UAS-mCD8-GFP* were crossed with *UAS-mCherry*. The progeny, *en-Gal4, UAS-mCD8-GFP/UAS-mCherry* female virgins were crossed with an *If/CyO* balancer stock. Larvae were screened under a Leica fluorescence scope. mCherry positive and GFP negative larvae were collected to build an *en-Gal4, UAS-mCherry* stock.

To create the *nub-Gal4, UAS-mCherry* driver, *nub-Gal4, UAS-mCherry; UAS-Nslmb-vhhGFP4/TM3, sb* was crossed with the *If/CyO* balancer stock, *nub-Gal4, UAS-mCherry/CyO; +/TM3, sb* and progeny were collected.

### Imaging of adult flies and processing of images

Female adult flies were kept at −20 °C for at least two hours before imaging. The flies were aligned on a small piece (5×5 cm) of printing paper. Images were taken under a dissection scope. For each fly, about twenty images of different focus were shoot by adjusting the scope. To get a deep focus image of a fly, the original images were processed using the Helicon Focus software (7.5.8, Helicon Soft Ltd). The images were imported into the Helicon Focus software. Method C (pyramid) was applied. The output image was saved in .tif format.

For taking fluorescent images, newly eclosed flies were pinched at the neck using a pair of forceps. The flies were allowed to recover for one hour. The images were taken under a Leica fluorescence scope and processed using the Helicon Focus software.

### Flp-out clone induction

To induce somatic clones, female virgins of *hs-Flp*^*122*^; *UAS-Flp*^*JD1*^*/CyO, Act-GFP*^*JMR1*^; *Act5C > CD2 > Gal4*^*S*^, *UAS-mCD8GFP*^*LL6*^*/TM6b* were crossed with *w*^*1118*^ and *UAS-bab2*, respectively. Embryos were collected in an 8-h time window. The newly hatched larvae were synchronized. A heat shock (37 °C in a water bath) was carried out 84 h after larval hatching to induce clones. The imaginal discs were dissected 34 hours after clone induction.

### Immunohistochemistry

*Drosophila* wing discs were dissected in phosphate-buffered saline pH 7.2 (PBS) and fixed in 4% paraformaldehyde (pH=7.4) for 30 min at room temperature, washed one time in PBS, three times in PBST (0.3% Triton-X 100 in PBS), and pre-incubated in PBST with 0.5% normal donkey goat serum. The specimens were incubated with primary antibodies overnight at 4 °C, then washed three times in PBST and incubated with secondary antibodies for two hours at room temperature, followed by washing four times in PBST and one time in PBS. The following antibodies were used: mouse anti-β-Galactosidase 1:100 (40-1a, Developmental Studies Hybridoma Bank, DSHB), rabbit anti-Cleaved Caspase 3 1:300 (Cell Signaling), rat anti-Bab2 1:10000 (Jean-Louis Couderc), Alexa Fluor 488 donkey anti-mouse 1:500 (Invitrogen), Alexa Fluor 594 donkey anti-mouse 1:500 (Invitrogen), Alexa Fluor 647 donkey anti-rabbit 1:500 (Invitrogen), Alexa Fluor 488 goat anti-rabbit 1:500 (Eugene).

### Confocal Image processing and data collection

Images were processed with Fiji software. To quantify the average fluorescence intensity, the region of interest in an image was selected using the ‘Freehand selections’ tool and subsequently analyzed using ‘Analyze’, then ‘measure’. To quantify the fluorescence profile, the region of interest in an image was selected using ‘Rectangle’ tool, and then analyzed using ‘Analyze’, ‘Plot Profile’.

### Molecular cloning

To generate a UAS vector for overexpression of a cDNA encoding an N-terminal Flag (DYKDDDDK)-tagged Bab2 protein, the *bab2CD*(Couderc et al., 2002) was first PCR-amplified, digested with *Eco*RI (ThermoFisher) and *Xba*I (ThermoFisher), and cloned into the pUAST vector using the same restriction enzymes. The plasmid was extracted using a Plasmid Midi Kit (Cat# 12143, QIAGEN) according to the manufacturer’s protocol.

### S2 cell transfection

The Gal4-UAS system was used for expression of Flag tagged Bric à brac Conserved Domain (*UAS-flag-bab2CD*) in S2 cells. Four hours before transfection, S2 cells (approximately 1 x 10^6^ cells/ml) were seeded in a 6 well plate (2 ml/well). The *pAct-Gal4* and *pUAST-bab2CD* plasmids were co-transfected using Effectene Transfection Reagent (QIAGEN) according to the manufacturer’s protocol. Sixteen hours after transfection, the culture medium was replaced with fresh medium. The cells were harvested 24 hours after transfection.

### Chromatin immunoprecipitation (ChIP)

Chip was performed according to an online protocol (https://www.abcam.com/protocols/cross-linking-chromatin-immunoprecipitation-x-chip-protocol) with minor changes. S2 cells, 3-4 x 10^7^,were harvested and washed in ice-cold PBS three times. The pellet was resuspended in 2.4 ml ChIP lysis buffer and incubated for 10 min on ice. The lysate was sonicated in an Ultrasonic Processor (Vibra-Cell™) for 5 min (10s pulse/10s interval, 50% power). Dynabeads Protein A (Cat# 10002D, Novex) and Protein G (Cat# 10004D, Novex) were used in the experiments. Separation of beads and buffer was performed on a magnetic separation stand.

### Quantitative PCR (qPCR)

Reactions for qPCR were carried out using SYBR Green (ThermoFisher) in a Rotor-Gene 6000 cycler (Qiagen). Genomic DNA was used as a standard. The final concentration of primers was 300 nM.

The following primers were used:

dpp-Ch-F: AAACCGGTGAAAACCACAGC

dpp-Ch-R: TTGCTTTCAAAGAGCACCGC

wg-Ch-F: TTTCCGTGATACGGAGAGCG

wg-Ch-R: TTTCCAGTTGCGTTGCGAAG

rp49-F: AACCGCGTTTACTGCGGCGA

rp49-R: AGAACGCAGGCGACCGTTGG

### Data plot and statistical analysis

Data plots and statistical analyses were performed with GraphPad Prism 8.0 software.

## Acknowledgments

We gratefully acknowledge the Bloomington Drosophila Stock Center (NIH P40OD018537) for fly stocks, the Vienna Drosophila Resource Center (VDRC, www.vdrc.at) for transgenic fly stocks, and the Developmental Studies Hybridoma Bank, created by the NICHD of the NIH and maintained at The University of Iowa, Department of Biology, Iowa City, IA for the monoclonal antibodies used in this study. We thank Prof. Jean-Louis Couderc (Clermont Universitè) and Prof. Leonard Rabinow (University of Paris-Sud) for providing fly strains and antibodies. We also thank the late Prof. Heinrich Reichert (Basel University) for support, Susanne Flister (Basel University) for technical assistance, and Mattias Mannervik and Ulrich Theopold for constructive comments during manuscript preparation.

## Author contributions

Y.Z. conceived the project, designed and performed experiments, analyzed data and performed statistical analyses, prepared figures and wrote the manuscript; J.D. designed and performed experiments, analyzed data and prepared figures; A.D. analyzed data and wrote the manuscript, S.B. and Y.E. conceived, funded, and supervised the project, analyzed data and wrote the manuscript.

